# Receptor tyrosine kinase inhibitors trametinib and axitinib effectively target the neoplastic component of giant cell tumor of bone in vitro

**DOI:** 10.64898/2026.10.06.756991

**Authors:** Veronika Knechtova, Michal Mahdal, Iva Staniczkova Zambo, Dusan Zoufaly, Olga Piskareva, Petr Chlapek, Jan Skoda, Jakub Neradil

**Affiliations:** Department of Experimental Biology, Faculty of Science, Masaryk University, 625 00 Brno, Czech Republic.; International Clinical Research Center, St. Anne’s University Hospital, 656 91 Brno, Czech Republic.; First Department of Orthopaedic Surgery, St. Anne’s University Hospital and Faculty of Medicine, Masaryk University, 656 91 Brno, Czech Republic.; First Department of Pathology, St. Anne’s University Hospital and Faculty of Medicine, Masaryk University, 656 91 Brno, Czech Republic.; Cancer Bioengineering Group, Department of Anatomy and Regenerative Medicine, RCSI University of Medicine and Health Sciences, Dublin, Ireland.; Tissue Engineering Research Group, Department of Anatomy and Regenerative Medicine, RCSI University of Medicine and Health Sciences, Dublin, Ireland.

**Keywords:** Giant cell tumor of bone, Neoplastic stromal cells, Receptor tyrosine kinase, Tyrosine kinase inhibitors, Targeted therapy, Trametinib, Axitinib

## Abstract

**Background:** Giant cell tumor of bone (GCTB) is an intermediate osteolytic neoplasm characterized by a high rate of local recurrence. The tumor comprises two main cell populations: neoplastic stromal cells and osteoclast-like giant cells. Current adjuvant therapies primarily target osteoclast-like giant cells but have limited activity against the neoplastic stromal cells that drive tumor growth and recurrence. Here, we sought to identify actionable vulnerabilities in GCTB stromal cells by profiling druggable receptor tyrosine kinases (RTKs) and associated signaling pathways and evaluating their pharmacological inhibition in patient-derived in vitro models.

**Methods:** RTK activation and downstream signaling were assessed using phospho-protein arrays and western blotting in therapy-naïve and denosumab-treated GCTB patient samples and in patient-derived GCTB cell lines, representing clinically relevant models of the neoplastic component. Tyrosine kinase inhibitors (TKIs) targeting activated pathways were evaluated in patient-derived GCTB cell lines cultured under both two-dimensional (2D) and three-dimensional (3D) conditions. Effects on cell viability, proliferation, cell cycle distribution, and cell death were assessed using MTT and XTT assays, flow cytometry, live-cell imaging, and confocal microscopy.

**Results:** Several activated RTKs, including EGFR, IGF-IR, InsR, PDGFRα, and PDGFRβ, together with downstream signaling proteins MEK1/2 and ERK1/2, were identified as potential therapeutic targets. Among the inhibitors tested, trametinib, a MEK1/2 inhibitor, and axitinib, a multi-target inhibitor of VEGFR1–3, PDGFRβ, and c-Kit, demonstrated the strongest antiproliferative and cytotoxic effects, markedly reducing cell viability and proliferation while promoting cell death in vitro. Importantly, trametinib showed selective activity against GCTB stromal cells compared with healthy controls, suggesting a potential therapeutic window.

**Conclusions:** This study identifies trametinib and axitinib as promising therapeutic candidates for targeting the neoplastic component of GCTB. Our findings provide a rationale for further preclinical and clinical evaluation of RTK-directed therapies to complement current treatments and specifically target the neoplastic GCTB stromal cells underlying tumor persistence and recurrence.

## 1. Introduction

Giant cell tumor of bone (GCTB) is an intermediate, locally aggressive, osteolytic neoplasm characterized by low metastatic potential but a high recurrence rate [1, 2]. GCTB accounts for approximately 5 % of all primary bone tumors and typically arises in adults between 20 and 45 years of age, predominantly affecting the meta-epiphyseal region of long bones [2]. Histologically, GCTB consists of two main cell types – neoplastic mononuclear stromal cells and osteoclast-like giant cells, the latter being responsible for the tumor’s osteolytic nature [3, 4]. The stromal cells represent the neoplastic component and, in more than 90% of cases, harbor a characteristic H3.3 G34W mutation [5]. These cells promote the formation of giant cells primarily through overexpression of receptor activator of nuclear factor kappa B ligand (RANKL), which induces the differentiation and fusion of monocytes into active multinucleated giant cells, resulting in excessive bone resorption [3, 6].

The primary treatment of choice is surgery; the choice of surgical approach and the use of adjunctive therapies depend on the tumor location and Campanacci stage [7]. Among adjuvant treatment options, denosumab—a human monoclonal antibody that binds RANKL, thereby inhibiting the formation and survival of osteoclast-like giant cells—is used particularly in cases of recurrent, unresectable, and Campanacci stage 3 GCTB [8]. Although denosumab treatment results in a complete depletion of giant cells, stromal cells are not eliminated and can drive recurrence after discontinuation of the treatment [9, 10, 1]. Therefore, there is a need for novel therapeutic strategies that combine denosumab with agents selectively targeting the neoplastic component of GCTB to reduce recurrence rates and improve patient outcomes.

The receptor tyrosine kinase (RTK) family comprises cell surface receptors involved in fundamental cellular signaling pathways, including those regulating cell growth, proliferation, differentiation, survival, migration, angiogenesis, and metabolism [11, 12]. Given their essential role in these processes, dysregulation of RTK signaling is a key driver of the initiation and progression of numerous cancers. This recognition has led to the development of receptor tyrosine kinase inhibitors (TKIs) that selectively target specific RTKs, many of which have received FDA approval for clinical use, with ongoing research focused on the development of novel and improved agents [11–13].

Several studies investigating novel therapeutic targets in GCTB stromal cells have revealed a distinct RTK activation profile in this neoplastic component [14–18]. We previously reported increased phosphorylation of specific RTKs and downstream signaling proteins after denosumab treatment, including epidermal growth factor receptor (EGFR), insulin-like growth factor 1 receptor (IGF-1R), platelet-derived growth factor receptor β (PDGFRβ), and ERK1/2 [14]. Furthermore, EGFR expression has been shown to positively correlate with GCTB progression and tumor recurrence [15, 18]. Importantly, the identified stromal cell-specific RTK activation patterns already show a strong translational potential. We previously demonstrated that treatment with a combination of denosumab and sunitinib, a PDGFRβ, vascular endothelial growth factor receptor 2 (VEGFR2), and c-Kit inhibitor, resulted in complete depletion of both stromal and giant cells in a patient with GCTB [14]. Moreover, a recent clinical study evaluating a neoadjuvant regimen combining denosumab and sunitinib in a small cohort of patients with GCTB reported that the treatment was well tolerated, induced complete depletion of giant cells and variable regression of stromal cells (mean regression, 62.8% ± 32.2%), resulting in 80% of patients remaining disease-free after 2 years [19]. Collectively, these findings suggest that concomitant targeting of RANKL and activated RTK signaling may represent a promising therapeutic strategy for patients with GCTB.

This study aimed to identify novel molecular targets in the neoplastic component for the treatment of GCTB. First, differences in RTK signaling profiles were analyzed in patient-derived samples following denosumab treatment and compared with those from untreated GCTB patients. Based on the observed RTK activation patterns, selected TKIs were evaluated for their therapeutic potential in GCTB cell lines under both 2D and 3D in vitro culture conditions. Our findings demonstrate that GCTB stromal cells are sensitive to several TKIs tested and reveal selective candidates that are more potent than sunitinib, suggesting their potential for targeting the neoplastic component of GCTB in combination treatment strategies.

## 2. Materials and methods

### 2.1. Tumor samples and cell lines

Tumor tissue samples were obtained from GCTB patients of the First Department of Orthopaedic Surgery, St. Anne’s University Hospital (Brno, Czech Republic). The Research Ethics Committee of St. Anne’s University Hospital approved the study protocol (approval no. 68V/2018), and written informed consent was obtained from each participant prior to their enrollment in this study. The samples included in this study (Tab. 1) were taken from 29 patients (17 males and 12 females; age range: 14–79 years old), some of the samples being from the same patient after recurrence. All patients included in this study were diagnosed with 2 or 3 Campanacci grade tumors. These patient tumor samples were also used to establish GCTB cell lines.

**Table 1:** Overview of the patientś samples included in this study.

| Sample number | Sex | Age at diagnosis | Localization | Campanacci Grade | Denosumab treatment (months) | Paired sample | Cell line | Western blot analysis | Phospho-protein analysis |
| --- | --- | --- | --- | --- | --- | --- | --- | --- | --- |
| 1 | M | 37 | Distal radius | 3 | - | - |  | x | x |
| 2 | M | 23 | Proximal fibula | 3 | - | 30 | GCTB6 | x | x |
| 3 | M | 42 | Distal femur | 2 | - | - |  | x | x |
| 4 | F | 44 | Proximal tibia | 2 | - | 9 | GCTB9 | x | x |
| 5 | F | 50 | Proximal tibia | 2 | - | - |  | x | x |
| 6 | M | 42 | Proximal femur | 3 | - | - |  | x | x |
| 7 | F | 18 | Distal ulna | 3 | - | 35 |  | x | x |
| 8 | F | 66 | Proximal<br>tibia | 2 | - | - |  | x |  |
| 9 | F | 46 | Proximal<br>tibia | 2 | - | 4 |  | x | x |
| 10 | M | 59 | Distal radius | 3 | - | 20 |  | x | x |
| 11 | F | 14 | metacarpus | 3 | - | 34 |  | x | x |
| 12 | M | 28 | Distal femur | 3 | - | - |  | x |  |
| 13 | M | 37 | Distal tibia | 2 | - | - |  | x |  |
| 14 | M | 33 | Proximal<br>tibia | 3 | 11 | - |  | x |  |
| 15 | M | 46 | Proximal<br>ulna | 3 | 6 | - |  | x | x |
| 16 | F | 15 | Proximal<br>fibula | 3 | 3 | - | GCTB10 | x | x |
| 17 | M | 40 | Distal radius | 2 | 3 | - |  | x | x |
| 18 | F | 29 | talus | 2 | 3 | - |  | x | x |
| 19 | F | 22 | sacrum | 3 | 18 | - |  | x | x |
| 20 | M | 60 | Distal radius | 3 | 3 | 10 |  | x | x |
| 21 | F | 50 | metacarpus | 3 | 3* | - |  | x |  |
| 22 | F | 61 | Proximal<br>tibia | 3 | 3 | - |  | x | x |
| 23 | M | 78 | Distal ulna | 3 | 3 | 25 |  | x |  |
| 24 | M | 25 | Distal ulna | 3 | 3 | - |  | x |  |
| 25 | M | 79 | Distal ulna | 3 | 3 | 23 |  | x |  |
| 26 | M | 44 | Proximal<br>tibia | 3 | - | 27 | GCTB40 |  | x |
| 27 | M | 45 | Proximal<br>tibia | 3 | 3* | 26 | GCTB46 |  |  |
| 28 | F | 18 | Distal ulna | 3 | 3* |  | GCTB47 |  |  |
| 29 | M | 27 | cuboideum | 3 | 3 |  |  |  | x |
| 30 | M | 26 | Proximal<br>fibula | 3 | 6 | 2 |  |  | x |
| 31 | M | 27 | Distal radius | 3 | 3 |  |  |  | x |
| 32 | M | 19 | Distal radius | 3 | - |  |  |  | x |
| 33 | F | 16 | Pollex-<br>metacarpus | 3 | - | 11 |  |  | x |
| 34 | F | 20 | Distal ulna | 3 | - | 7 |  |  | x |

\* plus 1 month treatment with sunitinib

### 2.2. Cell culture and treatment

The following cell lines were used in this study: (i) six in-house patient-derived GCTB cell lines established from tumor tissue with written informed consent (Tab. 1); (ii) human neonatal dermal fibroblasts NDF-2 (#CC-2509; Lonza Bioscience, Durham, NC, USA; a kind gift of Dr. Tomáš Bárta, Masaryk University, Brno). All cell lines were authenticated by STR profiling (Generi Biotech, Hradec Králové, Czech Republic) and routinely tested for mycoplasma contamination via PCR [20]. GCTB-derived cell lines were cultured in DMEM low glucose medium (#LM-D1100, Biosera, Nuaillé, France) supplemented with 20% fetal bovine serum (FBS) (#S-FBS-CO-015, Serana, Pessin, Germany), 2 mM L-glutamine (#XC-T1715, Biosera), 100 IU/ml penicillin and 100 μg/ml streptomycin (#XC-A4122, Biosera). NDF-2 cell line was cultured in DMEM high glucose (#LM-D1112, Biosera) supplemented with 10% FBS, 1% non-essential amino acids (#XC-E1154, Biosera), 2 mM L-glutamine, 100 IU/ml penicillin and 100 μg/ml streptomycin. All cell lines were cultured in a humidified 5% CO_2_ atmosphere at 37 °C.

The following low-molecular inhibitors targeting RTKs were used for treatment: sunitinib (#12328, Cell Signaling Technology, CST, Danvers, MA, USA), axitinib (#S1005, Selleckchem, Houston, TX, USA), trametinib (#GEN1590424, APEXBIO, Houston, TX, USA), crizotinib (#PZ0191, Sigma-Aldrich, St. Louis, MO, USA), and ceritinib (#S7083, Selleckchem).

### 2.3. MTT assay

Cells were seeded in 96-well plates (#30096, SPL Life Sciences, Gyeonggi-do, South Korea) at a density of 2000 cells/well and, 24 h after seeding, treated with the selected inhibitors for six days. After treatment, cell viability was measured using the MTT assay; thiazolyl blue tetrazolium bromide (#M2128, Sigma‒Aldrich) was added to a final concentration of 455 µg/ml. After 3 h of incubation under standard conditions, the medium was replaced with 200 µl of DMSO, and the absorbance was measured at 570 nm with a reference absorbance at 620 nm wavelength using a Sunrise absorbance reader (Tecan, Männedorf, Switzerland). The tested concentrations were normalized to untreated controls.

### 2.4. Growth analysis

Cells were seeded in 96-well plates at density 2000 cells/well, and 24 h after seeding, treated with the selected inhibitors for six days. Meanwhile, cell confluency was measured every 24 h using live cell imaging system Incucyte® SX1 (Sartorius, Göttingen, Germany) and plotted relative to initial values and to untreated control.

### 2.5. Flow cytometry

For all experiments, cells were harvested and enzymatically dissociated into single-cell suspension using Accutase (#LM-T1735, Biosera) and analyzed after staining using CytoFLEX S flow cytometer (Beckman Coulter, Brea, CA, USA). For cell viability, cells were suspended in PBS with 3% FBS and 2 mM EDTA (#ED2SS, Sigma-Aldrich) and incubated with 5 nM SYTOX™ Red Dead Cell Stain (#S34859, Invitrogen, Carlsbad, CA, USA) for 15 min on ice as per the manufacturer’s instructions.

For cell cycle analysis, the harvested cells were washed with PBS and then fixed with 70% ethanol for 10 min on ice. After fixation, cells were washed with PBS and resuspended in Vindelov’s solution (1% 1 M Tris (pH 8), 10-70 μg/ml RNase, 0.6 mg/ml NaCl, 50 μg/ml propidium iodide) diluted 1:1 with PBS, and incubated for 30 min at 37 °C.

### 2.6. Western blotting

GCTB tissue samples were manually dissected and lysed in LB17 lysis buffer (Proteome Profiler Human Phospho-RTK Array Kit; #ARY001B, RCD Systems, Minneapolis, MN, USA) supplemented with cOmplete™ Mini Protease Inhibitor Cocktail (#11836170001, Roche, Basel, Switzerland). In case of GCTB cell lines, whole-cell extracts were collected using RIPA lysis buffer (2 mM EDTA, 1% IGEPAL® CA-630, 0.5% SDS, 8.7 mg/ml sodium chloride, 5 mg/ml sodium deoxycholate, 50 mM Tris-HCl) supplemented with cOmplete™ Mini Protease Inhibitor Cocktail and PhosSTOP (#4906837001, Roche). 10 µg of total proteins were loaded onto 10% SDS-polyacrylamide gels, electrophoresed and blotted onto PVDF membranes (#1620177, Bio-Rad Laboratories, Hercules, CA, USA). The membranes were blocked with 5% not-fat dry milk or bovine serum albumin (#A7906, Sigma-Aldrich) in Tris-buffered saline with 0.05% Tween-20 (#93773, Sigma-Aldrich) for at least 1 h and incubated with primary antibodies overnight on rocking platform at 4°C. Subsequently, incubation with secondary HRP-linked antibody was carried out at RT for at least 1 h. Chemiluminescent detection was performed following a 5 min incubation with ECL™ Prime Western Blotting Detection Reagent (#RPN2236, Cytiva, Marlborough, MA, USA) using Azure C600 imaging system (Azure Biosystems, Dublin, CA, USA).

Densitometric analysis of western blotting images was performed using gel analysis tool in ImageJ (Fiji) software (NIH, Bethesda, MD, USA). A protein of interest was normalized to a loading control, GAPDH, detected on the same gel.

The following primary antibodies were used for immunodetection: anti-EGFR (1:1000; #2646, CST), anti-phospho-EGFR (Y1148) (1:500; #4404, CST), anti-PDGFRβ (1:1000; #3169, CST), anti-phospho-PDGFRβ (Y751) (1:1000; #4549, CST), anti-IGF-IRβ (1:1000; #9750, CST), anti-phospho-IGF-IRβ (Y1135/1136) (1:1000; #3024, CST), anti-Met (1:1000; #8198, CST), anti-phospho-Met (Y1234/1235) (1:1000; #3077, CST), anti-MEK1/2 (1:1000; #9122, CST), anti-ERK1/2 (1:1000; #4695, CST), anti-phospho-ERK1/2 (T202/Y204) (1:1000; #4370, CST), anti-HSP60 (1:1000; #12165, CST), anti-c-Jun (1:1000; #9165, CST), anti-H3.3 G34W (1:500; #31-1145-00, RevMAb

Biosciences USA, Inc, San Francisco, CA, USA), anti-H3 (1:1000; #14269, CST), anti-GAPDH (1:10 000; #sc-365062, Santa Cruz Biotechnology, Dallas, TX, USA). The following secondary antibodies were used: goat anti-rabbit IgG (1:5000, CST, #7074) and horse anti-mouse IgG (1:5000, CST, #7076).

### 2.7. Phosphoprotein array analysis

RTK phosphorylation was analyzed using the Proteome Profiler™ Human Phospho-RTK Array Kit (#ARY001B, RCD Systems) according to the manufacturer’s instructions. Tissue samples were lysed in Lysis Buffer 17, centrifuged (14,000 × g, 5 min), and 300 µg of total protein was applied per array. A detailed description of the method has been published previously [21]. Chemiluminescent signals were detected using the Azure C600 imaging system (Azure Biosystems, Dublin, CA, USA) or captured on an X-ray film and quantified using ImageJ. Raw pixel density values of individual spots were background-subtracted, and the duplicate spots corresponding to same protein were averaged. The resulting values were corrected by subtracting the values obtained from negative controls, in which Lysis Buffer 17 was used instead of the protein lysate.

### 2.8. Collagen-nHA scaffold fabrication and seeding

The scaffolds were manufactured as previously described by others [22, 23]. Briefly, collagen slurry (0.5% (w/v)) was fabricated by blending microfibrillar bovine tendon collagen I (Integra Life Sciences, Inc., Princeton, NJ, USA) with 0.05 M glacial acetic acid at 4 °C for several hours using an overhead blender (IKA Works, Inc., Wilmington, NC, USA). The nanohydroxyapatite (nHA) particles were synthesized by dissolving 2.27 g Na_3_PO_4_, 0.02 g NaOH, and 200 μl Darvan in 100 ml of dH_2_O, then slowly combining the solution with a solution of 1.47 g CaCl_2_ in 100 ml of dH_2_O. The resulting nHA suspension was centrifuged at max speed for 60 min, supernatant was discarded and nHA was resuspended in 60 ml of dH2O. The resuspended nHA was mixed with the collagen slurry during the blending process, resulting in nHA:collagen ratio 2:1. The slurry was degassed using vacuum pump, poured into stainless steel molds, and freeze-dried by cooling to -40 °C at 0.9 °C/min, followed by sublimation step for 17 h at 0 °C and 200 mTorr (Martin Christ Epsilon 2-4 LSC Plus, Martin Christ, Osterode am Harz, Germany). Freeze-dried scaffolds were dehydrothermally crosslinked in a vacuum oven (VacuCell, MMM, Munich, Germany) at 50 mTorr, 105 °C for 24h. Afterwards, scaffolds were rehydrated in PBS, chemically crosslinked with 14 mM N-(3-Dimethylamino propyl)-N’-ethylcarbodiimide hydrochloride (#E7750, Sigma-Aldrich) and 5.5 mM N-hydroxysuccinimide (#130672, Sigma-Aldrich) in dH2O, and sterilized in 70% ethanol for 30 min. Scaffolds were placed in an ultra-low-attachment 24-well plate (#CLS3473, Corning, Corning, NY, USA) (1 scaffold per well), and 1×10^5^ cells in 10 μl of media were dropped on one side of the scaffold. After 30 min of incubation, 1.5 ml of cell culture media was added to each well.

### 2.6. XTT assay

XTT assay was performed 4 days after seeding cells on scaffolds (day of treatment) and again after 6 days of treatment. Cell culture media was discarded, and scaffolds were rinsed with PBS. 1 ml of fresh complete medium without phenol red was added (#11880028, Gibco, Grand Island, NY, USA), followed by 500 μl of 1 mg/ml 2,3-Bis(2-methoxy-4-nitro-5-sulfophenyl)-2H-tetrazolium-5-carboxanilide (#X4626, Sigma-Aldrich) with 7.5 µg/ml Phenazine methosulfate (#P9625, Sigma-Aldrich). After 5 h of incubation under standard conditions, the medium was transferred into 96-well plate and the absorbance was measured at 450 nm with a reference absorbance at 620 nm wavelength using a Sunrise absorbance reader (Tecan, Männedorf, Switzerland). The values after treatment were normalized to initial values and tested inhibitors were normalized to untreated controls.

### 2.10. Live/dead staining

Staining solution was prepared by diluting 8 μg/ml of fluorescein diacetate (FDA) (#F7378, Sigma-Aldrich) and 5 μM of TO-PRO^TM^-3 Iodide (642/661) (#T3605, Invitrogen) in cell culture media without FBS. Scaffolds with seeded cells were rinsed in PBS and incubated in staining solution for 30 min at RT in dark. Stained scaffolds were analyzed with a confocal microscope (Leica SP8, Wetzlar, Germany).

### 2.11. Histological staining

Cells on collagen-nHA scaffolds were fixed in 4% paraformaldehyde for 30 min and embedded in paraffin to obtain FFPE blocks. Sections of the samples were stained with hematoxylin-eosin using an automated Medite TST 44 slide stainer (Medite Medical, Burgdorf, Germany). Furthermore, samples were stained with anti-H3.3 G34W (1:200; #31-1145-00, RevMAb) and mounted on a glass slide.

### 2.12. Statistical analysis

All the statistical analyses were performed with at least three biological replicates (details in the figure legends) using GraphPad Prism 8.0.2 software. Mann-Whitney U test was performed for comparison between two groups. One-way ANOVA with Welch’s correction followed by post hoc Dunnett’s test, one-way ANOVA with Dunnett’s multiple comparisons test, Brown-Forsythe and Welch ANOVA test with Dunnett’s T3 multiple comparisons test, RM two-way ANOVA with the

Geisser-Greenhouse correction and Dunnett’s multiple comparisons test, or Kruskal-Wallis with Dunńs multiple comparisons test were used to compare multiple groups. p values < 0.05 were considered statistically significant; *p<0.05, **p<0.01, ***p<0.001, #p<0.0001.

## 3. Results

### 3.1. Giant cell depletion following denosumab treatment is associated with a distinct RTK signaling in GCTB

To investigate RTK expression profiles in GCTB and assess their potential as therapeutic targets, we first analyzed the expression of selected RTKs and downstream signaling proteins in patient tumor samples (Fig. 1). The cohort included 12 samples obtained after denosumab treatment and 13 samples from patients who underwent surgery without prior denosumab therapy. As reported previously [4, 14], histological evaluation revealed that all denosumab-treated samples contained exclusively neoplastic stromal cells, whereas untreated samples comprised both neoplastic stromal cells and osteoclast-like giant cells (Fig. S1). Distinct RTK expression patterns were observed between the two groups. The downstream signaling proteins c-Jun and HSP60 were significantly more abundant in untreated GCTB tissue, suggesting that their expression is associated with the presence of giant cells. In contrast, RTK PDGFRβ was highly expressed, and the signaling proteins MEK1/2 and ERK1/2 exhibited high levels of phosphorylation in denosumab-treated samples, indicating predominant expression in neoplastic stromal cells. To determine whether expressed RTKs also exhibit elevated phosphorylation and are thus activated, we performed phospho-protein array analyses.

**Fig. 1.**
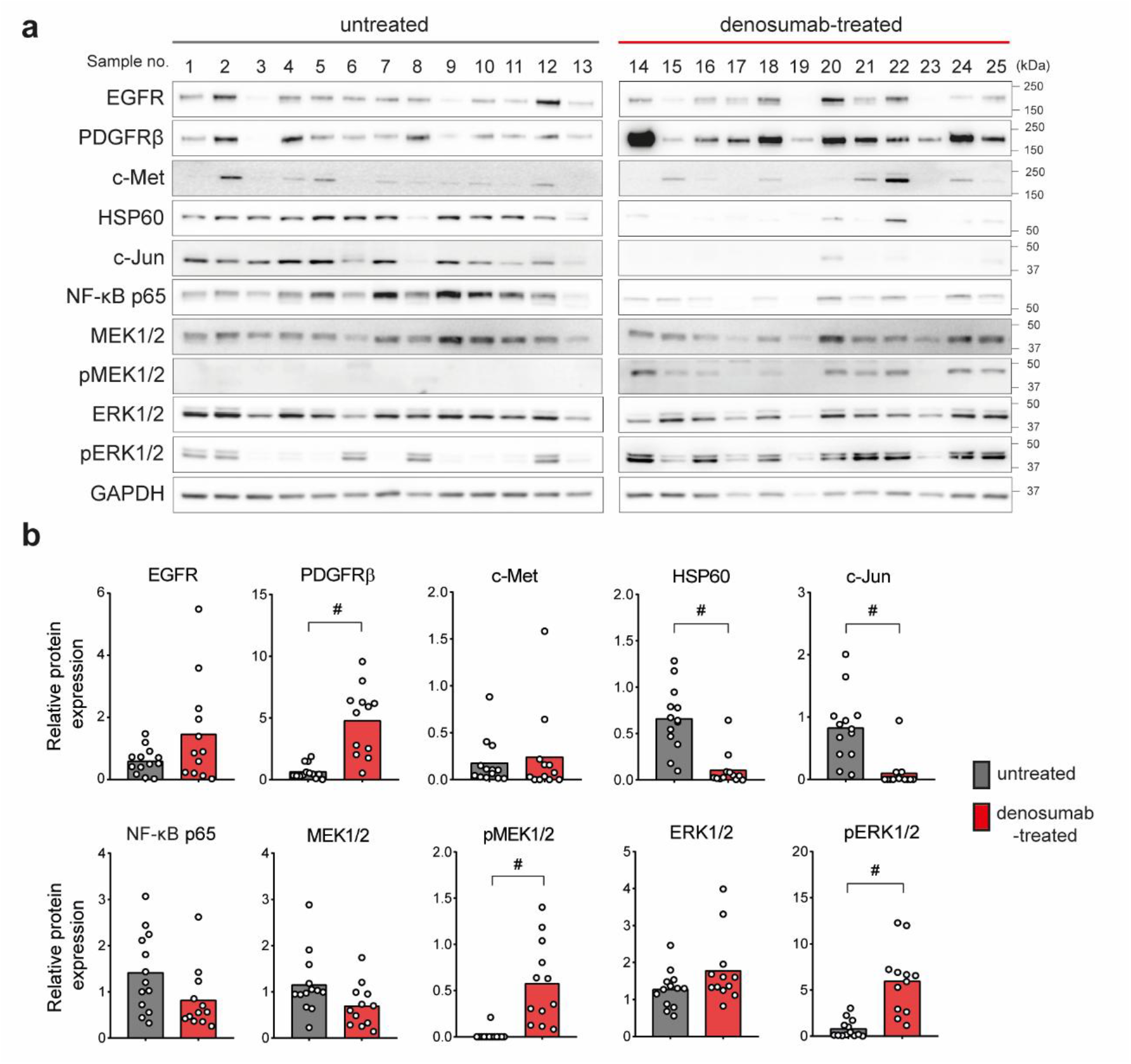
Denosumab treatment profoundly alters the expression of RTKs and downstream signaling proteins in GCTB tissue. **(a)** Western blot analysis of RTKs and downstream signaling proteins in 13 untreated and 12 denosumab-treated GCTB patient samples and **(b)** densitometric analysis. The values were normalized to a calibrator sample used on all membranes. Statistical significance was determined by Mann-Whitney U test, #p<0.0001.

### 3.2. Phosphorylation of expressed RTKs confirms their activation in denosumab-treated GCTB patient samples

To assess RTK phosphorylation levels in patient tumor samples, we performed phospho-protein array analysis (Fig. 2, S2). A total of 10 samples obtained from denosumab-treated patients and 14 samples from treatment-naïve patients were included in the analysis. Unsupervised hierarchical clustering grouped the majority of denosumab-treated samples into a distinct cluster. Two untreated samples clustered together with treated cases, suggesting that a subset of untreated tumors may share a similar RTK activation profile. Samples from treatment-naïve patients exhibited increased phosphorylation of ROR2 and c-Ret. In contrast, samples obtained following denosumab treatment showed significantly increased phosphorylation of EGFR, IGF-IR, InsR, PDGFRα, and PDGFRβ. In the case of PDGFRβ, the elevated phosphorylation levels were consistent with increased total protein expression. This distinct shift in RTK phosphorylation patterns following denosumab treatment may reflect either a direct effect of denosumab on GCTB stromal cells or alterations resulting from giant cell depletion and the consequent disruption of signaling between these two cellular populations. Regardless of the underlying mechanism, RTKs exhibiting high expression and phosphorylation levels in stromal cells following denosumab treatment may represent promising therapeutic targets.

**Fig. 2.**
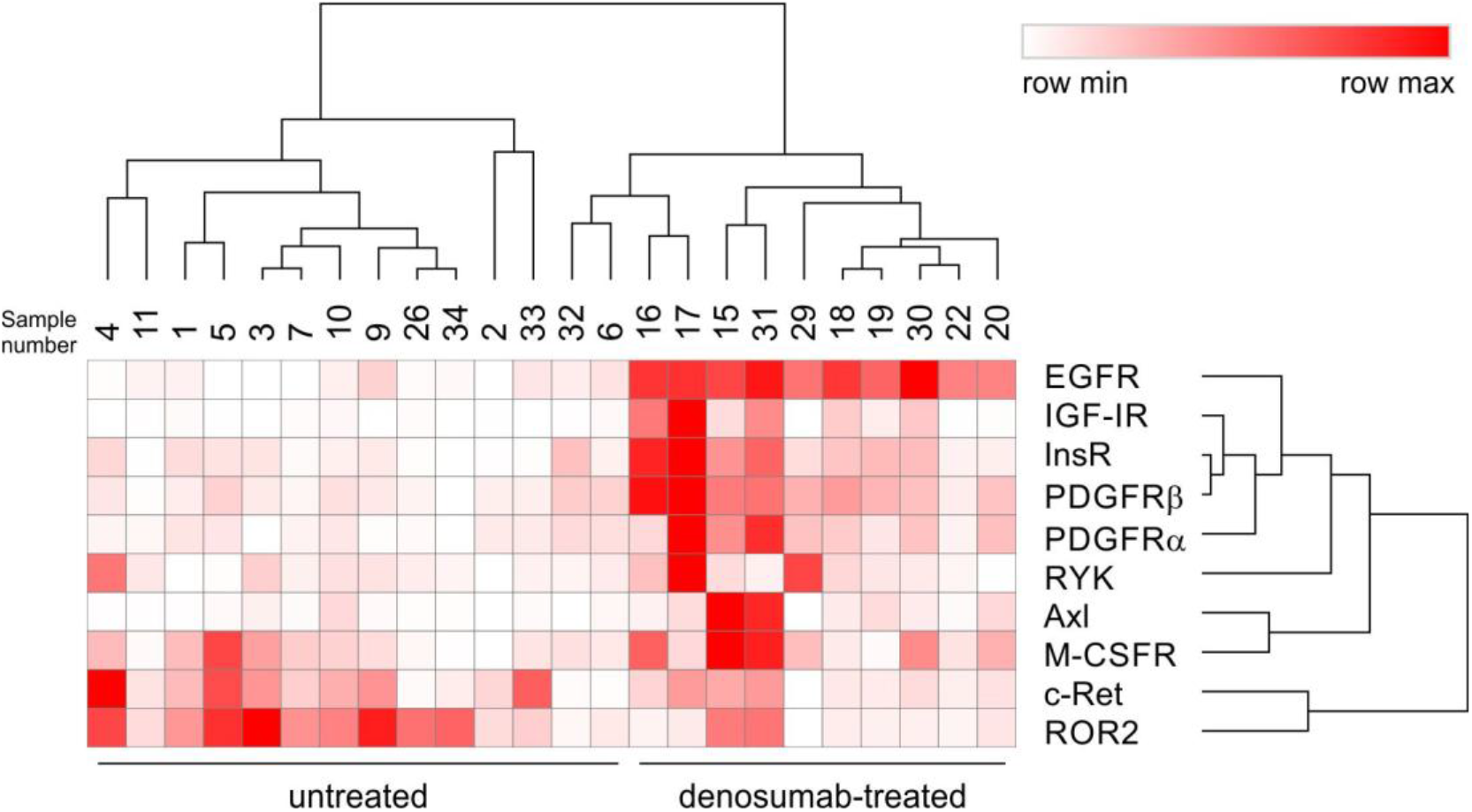
Giant-cell depletion by denosumab treatment reshapes RTK phosphorylation profiles in GCTB tissues. 10 denosumab-treated and 14 samples from GCTB patients without prior denosumab treatment were analyzed using phospho-RTK arrays. Obtained data were visualized as a heatmap and analyzed using hierarchical clustering. Hierarchical clustering was performed using the 10 most variable RTKs across all samples. Data were clustered using one minus Pearson correlation and average linkage.

### 3.3. Patient-derived GCTB cell lines present similar RTK profiles as denosumab-treated tumor samples, providing a relevant in vitro model

To establish an *in vitro* model of the neoplastic component of GCTB, we derived six cell lines from patient tumor tissues. As osteoclast-like giant cells do not propagate under standard cell culture conditions, they are progressively lost during passaging, resulting in cultures composed exclusively of stromal cells. Consequently, GCTB cell lines are expected to exhibit RTK expression profiles comparable to those of denosumab-treated GCTB samples, in which osteoclast-like giant cells have been depleted. The presence of the characteristic histone H3.3 G34W mutation detected in all derived cell lines validated their GCTB stromal cell identity (Fig. 3a). To assess the suitability of these patient-derived GCTB cell lines as an in vitro model for TKI testing, we analyzed RTK expression and phosphorylation. Western blot analysis demonstrated RTK expression across all cell lines, with some heterogeneity among samples (Fig. 3b and 3c). The expression profiles of stromal cell lines were compared with those of healthy human fibroblasts (NDF-2 cell line). These fibroblasts served as healthy control throughout the study, due to their shared mesenchymal origin with GCTB stromal cells, enabling assessment of both the specificity and potential toxicity of the tested compounds.

**Fig. 3.**
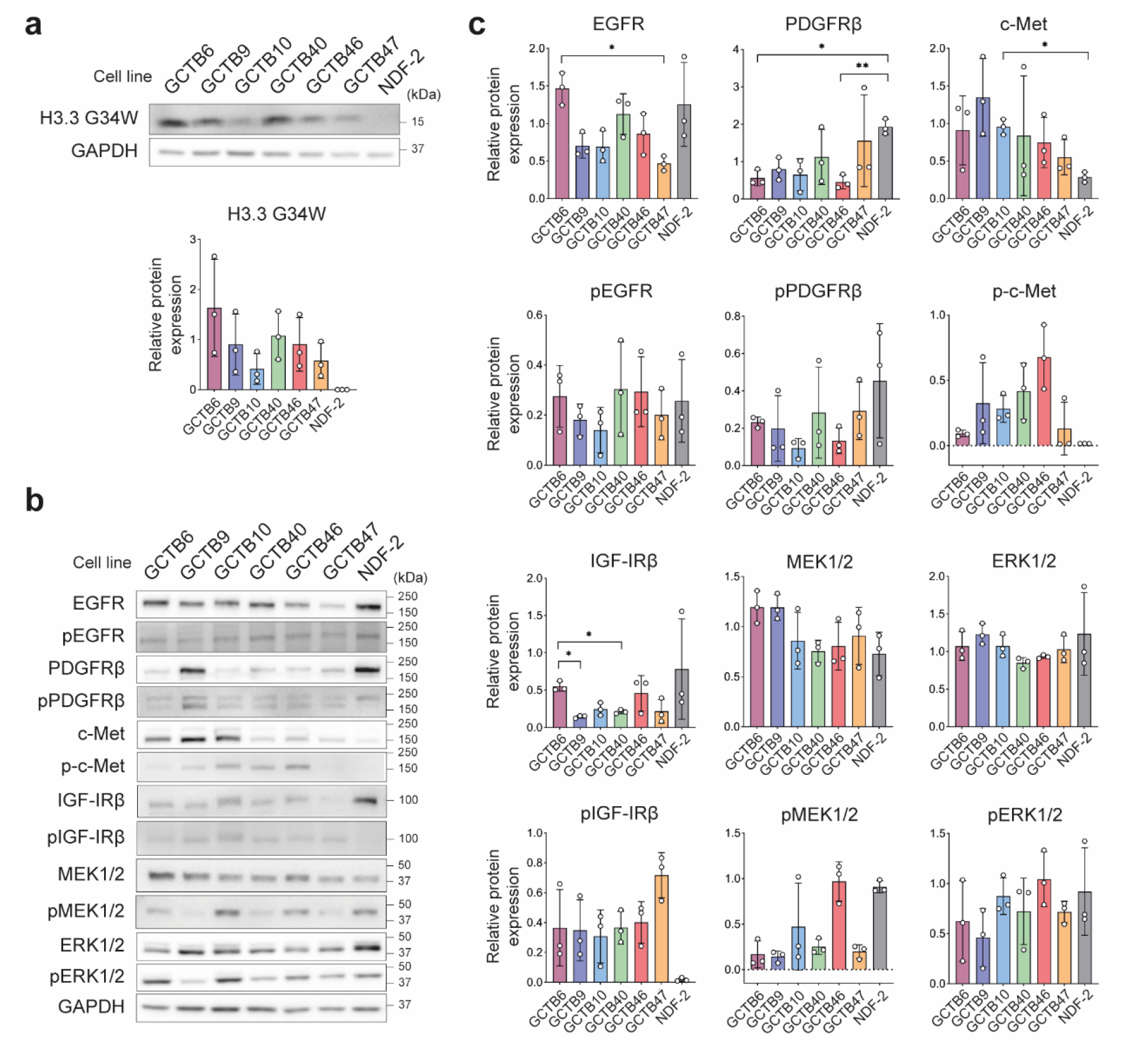
Patient-derived GCTB cell lines represent a relevant model of neoplastic stromal cells, reflecting the RTK signaling landscape of denosumab-treated GCTB. **(a)** Western blot and densitometric analysis of GCTB stromal cells-specific H3.3 G34W mutation. **(b,c)** Western blot (b) and densitometric analysis (c) of the expression and phosphorylation of certain RTKs and downstream signaling proteins in 6 patient-derived GCTB cell lines and NDF-2 cell line. Data are presented as mean ± SD, biological n = 3. Statistical significance was determined by one-way ANOVA with Welch’s correction followed by post hoc Dunnett’s test, *p<0.05, **p<0.01.

### 3.4. Patient-derived GCTB stromal cells are sensitive to specific tyrosine kinase inhibitors

Based on our findings and prior reports, we hypothesized that inhibitors targeting highly expressed and phosphorylated RTKs or their downstream signaling proteins could directly affect GCTB stromal cells, providing a promising therapeutic strategy. Previous studies demonstrated that sunitinib, a PDGFRβ and VEGFR2 inhibitor, significantly reduced the viability of GCTB stromal cells *in vitro* [14] and, in clinical settings, induced tumor regression, symptomatic improvement, and reduction in the size of metastases in patients with GCTB [14, 24]. Furthermore, a recent clinical study evaluating a neoadjuvant regimen combining denosumab and sunitinib in a small cohort of patients with GCTB reported substantial but variable stromal cell regression (mean, 62.8% ± 32.2%) and a reduced local recurrence rate of 12.5% [19]. Despite these promising results, the pronounced variability in response to sunitinib across patients prompted us to evaluate additional TKIs as potential therapeutic options.

Based on the RTK expression and phosphorylation patterns observed in GCTB tumor samples following denosumab treatment, we selected several small-molecule inhibitors targeting these receptors or their downstream signaling pathways: axitinib (PDGFRβ and VEGFR1/2/3 inhibitor), trametinib (MEK1/2 inhibitor), ceritinib (ALK, IGF-1R, and InsR inhibitor), and crizotinib (ALK and c-Met inhibitor). Their effects on the viability and proliferation of patient-derived GCTB cell lines were assessed *in vitro* and compared with those of the current leading agent, sunitinib. To evaluate specificity and potential toxicity, the same inhibitors were also tested on healthy human fibroblasts (NDF-2 cell line).

First, we analyzed the effects of each inhibitor on cell viability using MTT assay (Fig. 4a) and live-cell imaging (Fig. 4b). As treatment responses showed minimal variability among the six patient-derived GCTB cell lines tested, the data were pooled and compared with those of the NDF-2 cell line. MTT assays revealed that only the highest concentration of sunitinib (10 μM) substantially reduced GCTB cell viability to approximately 20% of control levels, whereas lower concentrations reduced viability to approximately 80% of control. In contrast, trametinib treatment led to a pronounced reduction in viability at concentrations ranging from 10 to 0.1 μM. Similarly, axitinib reduced viability to approximately 20% at 10 μM and to approximately 40% at 1 μM. Overall, axitinib and trametinib reduced GCTB stromal cell viability more effectively than sunitinib in a dose-dependent manner while exhibiting lower toxicity toward fibroblasts. In contrast, while 10 μM ceritinib or crizotinib reduced GCTB cell viability to nearly 0%, they also showed equivalent effects in fibroblasts, suggesting nonspecific cytotoxicity or off-target effects. Notably, live-cell imaging demonstrated a similar decrease in relative growth rates in both GCTB and NDF-2 cells following inhibitor treatment, indicating proliferative arrest across all cell lines (Fig. 4b).

**Fig. 4.**
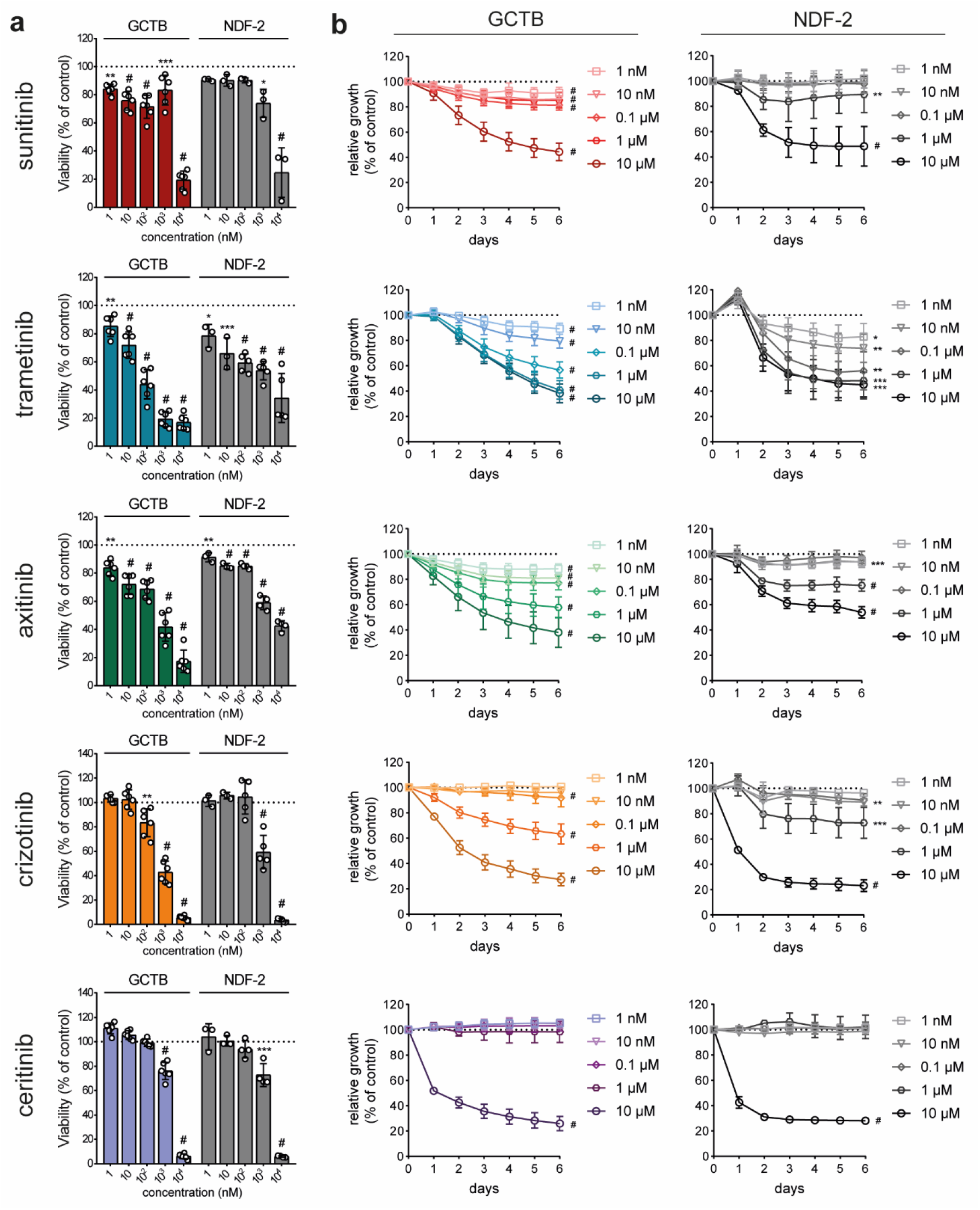
Targeted inhibition of specific RTKs reduces viability and inhibits proliferation across a panel of GCTB cell lines. **(a)** MTT assay cell viability analysis of 6 pooled GCTB cell lines (GCTB6, GCTB9, GCTB10, GCTB40, GCTB46, GCTB47) and neonatal dermal fibroblasts (NDF-2) after 6-day treatment with tyrosine kinase inhibitors sunitinib, axitinib, trametinib, crizotinib, and ceritinib. Data are presented as mean ± SD, biological n ≥ 3, technical n = 5. Statistical significance was determined by one-way ANOVA with Dunnett’s multiple comparisons test. **(b)** Live-cell imaging growth rate analysis of 6 pooled GCTB cell lines and NDF-2 cell line treated with indicated concentrations of tyrosine kinase inhibitors. Data are presented as mean ± SD, biological n = 3, technical n = 5. Statistical significance was determined by RM two-way ANOVA with the Geisser-Greenhouse correction and Dunnett’s multiple comparisons test, *p<0.05, **p<0.01, ***p<0.001, #p<0.0001.

To determine whether the observed effects were primarily cytostatic or cytotoxic, we quantified cell death by flow cytometry. Four GCTB cell lines were analyzed, and the data were pooled and compared with those from NDF-2 cells (Fig. 5a, S3a). Flow cytometric analysis demonstrated increased cell death in GCTB cell lines following treatment with axitinib and trametinib, confirming their greater efficacy relative to sunitinib. Overall, more pronounced reduction in metabolic activity and increased cell death observed in GCTB cells suggest greater sensitivity to treatment compared to NDF-2 cell line.

**Fig. 5.**
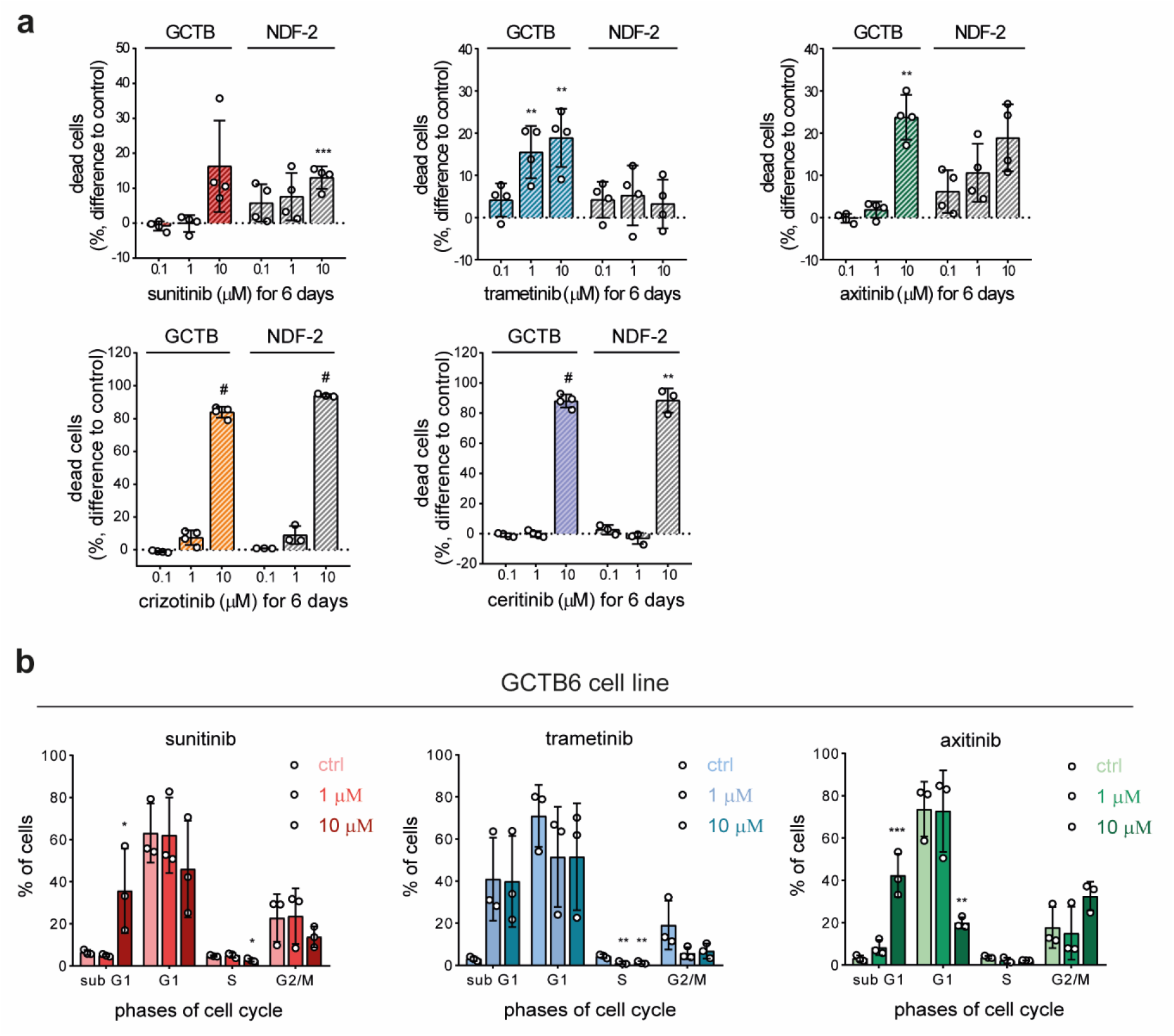
Targeted inhibition of specific RTKs induces cell death across a panel of GCTB cell lines. **(a)** Cell death rate of 4 pooled GCTB cell lines (GCTB6, GCTB10, GCTB40, and GCTB46) and NDF-2 cell line was measured after 6-day treatment with tyrosine kinase inhibitors (sunitinib, axitinib, trametinib, crizotinib, and ceritinib) by flow cytometry using SYTOX Red staining. Data are presented as the difference in percentages of SYTOX Red-positive dead cells after indicated treatment vs. respective untreated controls, biological n ≥ 3. Statistical significance was determined by Brown-Forsythe and Welch ANOVA test with Dunnett’s T3 multiple comparisons. **(b)** Changes in cell cycle in GCTB6 cell line after 6-day treatment with sunitinib, trametinib, or axitinib were evaluated using flow cytometry. Data are presented as mean ± SD, biological n = 3. Statistical significance was determined by ordinary one-way ANOVA with Dunnett’s multiple comparisons test, *p<0.05, **p<0.01, ***p<0.001, #p<0.0001.

Additionally, treatment-induced changes in cell cycle distribution were examined in the GCTB6 cell line after exposure to sunitinib, trametinib, and axitinib using flow cytometry (Fig. 5b, S3b). Consistent with the observed reduction in viability and increase in cell death, cell cycle analysis revealed an elevated sub-G1 population in GCTB6 cells treated with 10 µM sunitinib or axitinib and with 1 or 10 µM trametinib, even though not statistically significant.

Taken together, the combined viability, proliferation, cell death, and cell cycle analyses suggest that trametinib, followed by axitinib, is more effective than sunitinib in reducing GCTB stromal cell survival *in vitro*.3.5. Sunitinib, trametinib, and axitinib reduce viability and induce cell death in a 3D GCTB *in vitro* model

Although 2D cell culture remains the most widely used in vitro model in cancer research due to its low cost and high reproducibility, it is associated with several important limitations. Specifically, 2D culture represents an oversimplified system that lacks tissue complexity, spatial organization, and key components of the tumor microenvironment, including cell–cell interactions and gradients of nutrients, oxygen, and signaling molecules [25, 26]. The growing body of evidence highlights that cancer cells grown in 3D respond to drug treatment at clinically relevant doses [22, 27]. Hence, we evaluated selected TKIs in a 3D in vitro GCTB model using collagen–nHA scaffolds described earlier [22, 28]. Collagen–nHA scaffolds mimic the bone microenvironment and provide mechanical support for cell growth in 3D [23]. The ability of GCTB6 cells to infiltrate and survive within the scaffolds was confirmed by hematoxylin–eosin staining and immunohistochemical detection of the H3.3 G34W mutation (Fig. 6a). To evaluate the effects of the selected TKIs under 3D culture conditions, cells were seeded onto the scaffolds and, after 4 days of cultivation, treated with 1 or 10 µM of sunitinib, trametinib, or axitinib for 6 days. Cell viability was subsequently assessed using the XTT assay (Fig. 6b) and live/dead staining followed by confocal microscopy (Fig. 6c, d). All inhibitors showed comparable efficacy at 10 µM, reducing cell viability to below 40% of control levels and inducing substantial cell death. Importantly, trametinib exhibited activity even at 1 µM concentration, decreasing cell viability to approximately 60% and inducing cell death, whereas similar effects were not observed with sunitinib or axitinib. Notably, cells cultured under 3D conditions exhibited lower sensitivity to the inhibitors compared with 2D monolayer cultures. Even though considerable variability was observed among individual replicates, potentially due to differences in oxygen and nutrients availability within the scaffolds, or the extent of cellular infiltration into the scaffolds, which may have affected inhibitor penetration and efficacy, the overall trend of reduced GCTB cell viability and increased cell death following trametinib treatment remained evident.

**Fig. 6.**
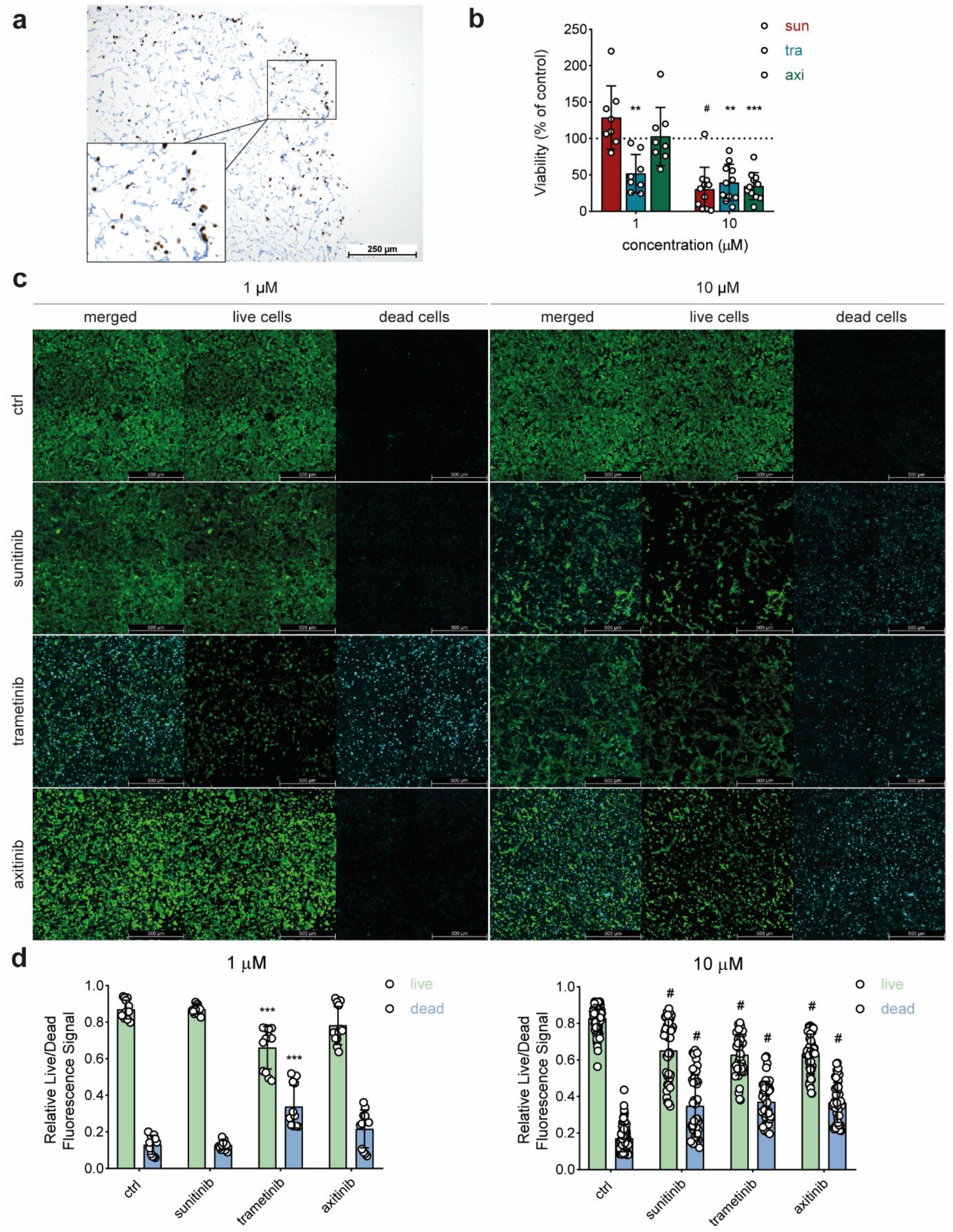
Targeted inhibition of specific RTKs decreases viability and induces cell death in GCTB cell line in 3D conditions. **(a)** A representative histological image of GCTB cells positive for H3.3 G34W mutation, on a collagen-nHA scaffold 4 days after seeding. Bar, 250 µm. **(b)** Cell viability analysis of GCTB cells on collagen-nHA scaffolds after 6-day treatment with 1 and 10 μM of TKI (sunitinib, trametinib, axitinib), measured by XTT assay. The values after treatment were normalized to values before treatment and tested inhibitors were normalized to untreated controls. Data are presented as mean ± SD, biological n ≥ 8. Statistical significance was determined by Kruskal-Wallis with Dunńs multiple comparisons test. **(c)** Representative confocal images of GCTB cells on collagen-nHA scaffold treated for 6 days with 1 and 10 µM of sunitinib, trametinib, or axitinib, stained with live (FDA)/dead (TO-PRO-3) staining. The displayed images show a maximal projection of z-stack. Bar, 500 µm. **(d)** Image analysis of the live/dead staining confocal images. The values were calculated as a raw integrated density of the live or dead cells divided by the sum of raw integrated density of both live and dead cells. Data are presented as mean ± SD, biological n ≥ 3, technical n = 4. Statistical significance was determined by Brown-Forsythe and Welch ANOVA with Tamhanés T2 multiple comparisons test, **p<0.01, ***p<0.001, #p<0.0001.

## 4. Discussion

GCTB is typically a locally aggressive tumor with low metastatic potential yet high recurrence rate, thus remaining a considerable clinical challenge. Surgery remains the primary treatment option and is, in selected cases such as unresectable, recurrent, or Campanacci stage 3 GCTB, combined with adjuvant denosumab therapy. Although denosumab effectively depletes giant cells, it does not eliminate neoplastic stromal cells, which may drive tumor recurrence following treatment discontinuation. Therefore, it is essential to develop novel therapeutic strategies combining denosumab with drugs selectively targeting neoplastic stromal cells.

In our previous study [14], we identified activated RTK signaling in GCTB stromal cells and hypothesized that TKIs targeting these activated RTKs could represent a viable therapeutic strategy. Supporting this hypothesis, a recent clinical study evaluated a neoadjuvant regimen combining denosumab with the multi-targeted TKI sunitinib, an inhibitor of VEGFR2 and PDGFRβ, in a small cohort of GCTB patients and demonstrated significant regression of both giant and stromal cells [19]. Similarly, the combination of denosumab with lenvatinib, a multikinase inhibitor targeting VEGFR1-3, FGFR1-4, PDGFRα, and KIT, exhibited moderate cytotoxicity in a GCTB cell line in vitro [29]. In addition, treatment of a patient with pulmonary metastatic GCTB using denosumab and the VEGFR2 inhibitor apatinib was associated with symptomatic improvement and tumor regression [16]. Although these findings suggest that combining denosumab with TKIs may represent a promising therapeutic strategy, the available evidence remains limited, and the reported responses of stromal cells have been variable. Therefore, we sought to evaluate additional TKIs as potential candidates for this combinatorial therapeutic approach.

First, we analyzed RTK expression and phosphorylation profiles in samples obtained from untreated and denosumab-treated GCTB patients. Samples collected after denosumab treatment exhibited a distinct RTK activation pattern, characterized by increased phosphorylation of EGFR, IGF-IR, InsR, MEK1/2, and ERK1/2, as well as both increased expression and phosphorylation of PDGFRβ. Based on these findings, we selected several TKIs for further evaluation, including axitinib (a VEGFR1/2/3 and PDGFRβ inhibitor), trametinib (a MEK1/2 inhibitor), ceritinib (an ALK, IGF-1R, and InsR inhibitor), and crizotinib (an ALK and c-Met inhibitor). Their effects were assessed in patient-derived GCTB cell lines cultured under both 2D and 3D conditions and compared with those of sunitinib.

Under 2D culture conditions, axitinib and trametinib reduced stromal cell viability and induced cell death more effectively than sunitinib in a dose-dependent manner while demonstrating lower toxicity toward healthy fibroblasts. In contrast, ceritinib and crizotinib exhibited marked toxicity, reducing the viability of both fibroblasts and GCTB cells to nearly 0%. In 3D culture conditions, trametinib demonstrated greater potency than both sunitinib and axitinib at lower concentrations (1 µM). At the highest tested concentration (10 µM), all evaluated inhibitors (sunitinib, axitinib, and trametinib) showed comparable efficacy, reducing cell viability to below 40% of control levels and inducing substantial cell death. Consistent with previous reports investigating drug sensitivity in 3D culture models [22, 25, 26, 30, 31], cells cultured on collagen-nHA scaffolds exhibited lower sensitivity to the inhibitors compared with 2D monolayer cultures. This reduced sensitivity may result from cell–ECM interactions, altered cellular phenotype and proliferation state, or shifts in cell signaling and gene expression profiles. Results obtained under 3D conditions also demonstrated greater variability between replicates, potentially reflecting nutrient and oxygen gradients, or the extent of cellular infiltration into the scaffolds. This greater variability highlights a limitation of scaffold-based assays, nevertheless, the overall trend of reduced cell viability and increased cell death following treatment remained evident. Since establishing animal models for GCTB has proven challenging, in vitro 3D models might provide a more physiologically relevant intermediate model, particularly for evaluating drug responses in a more tissue-like environment and improving the translation of results into the clinic.

Taken together, our findings identify trametinib and axitinib as promising therapeutic candidates against GCTB stromal cells and support further investigation of RTK/MAPK-targeted treatment strategies to prevent tumor recurrence and improve patient outcomes. Patient-derived GCTB cell lines showed greater sensitivity to trametinib and comparable sensitivity to axitinib relative to sunitinib under both 2D and 3D in vitro conditions. These results suggest that trametinib and axitinib could serve as alternatives to sunitinib in neoadjuvant regimens combined with denosumab, particularly in patients with limited stromal cell regression after denosumab and sunitinib treatment. Importantly, as both trametinib and axitinib are already FDA-approved and clinically used for other malignancies, they represent promising candidates for drug repurposing in GCTB treatment. However, these findings are preliminary and based solely on *in vitro* experiments using patient-derived GCTB cell lines; therefore, they cannot reproduce the complex interactions between stromal cells, osteoclast-like giant cells, immune cells, endothelial cells, and the extracellular matrix present in tumors. Even though the 3D collagen-nHA model provides a more tissue-like environment than 2D culture, it remains an incomplete representation of the GCTB microenvironment. Future studies should establish the mechanistic basis of the observed signaling changes, assess pharmacologically relevant drug concentrations, and validate efficacy and safety in appropriate preclinical models and clinical settings.

## Conclusions

In this study, we demonstrated that trametinib and axitinib inhibit proliferation, reduce viability, and induce cell death in GCTB stromal cells under both 2D and 3D in vitro culture conditions. Relative to sunitinib, GCTB cell lines exhibited greater sensitivity to trametinib and comparable sensitivity to axitinib. Collectively, these findings identify trametinib and axitinib as promising candidates for further preclinical investigation and potential clinical evaluation in GCTB, while providing additional support for targeting RTK-driven signaling pathways in the neoplastic stromal cells as a therapeutic strategy for GCTB.

## List of abbreviations

2D: two-dimensional
3D: three-dimensional
ECM: Extracellular matrix
EGFR: Epidermal growth factor receptor
GCTB: Giant cell tumor of bone
IGF-1R: Insulin-like growth factor 1 receptor
nHA: Nanohydroxyapatite
PDGFRβ: Platelet-derived growth factor receptor β
RANKL: Receptor activator of nuclear factor kappa B ligand
RTK: Receptor tyrosine kinase
TKI: Tyrosine kinase inhibitor
VEGFR2: Vascular endothelial growth factor receptor 2

## Declarations

### Ethics approval and consent to participate

This study was conducted according to the guidelines of the Declaration of Helsinki and approved by the Research Ethics Committee of St. Anne’s University Hospital (Brno, Czech Republic) (Protocol Code: 68V/2018, date of approval: 14 August 2018). All human tumor tissues and noncommercial patient-derived cell lines were obtained with written informed consent from patients or their legal guardians.

### Consent for publication

Written informed consent for publication of the clinical data was obtained from all patients or their legal guardians.

### Availability of data and materials

The datasets used and/or analysed during the current study are available from the corresponding author on reasonable request.

### Authors’ contributions

VK performed most of the experiments, analyzed the data, and wrote the manuscript. MM provided the clinical samples and patient data. ISZ, and DZ conducted the experiments and acquired data related to immunohistochemical analysis. OP provided methodology and facilities for manufacturing the scaffolds and edited the manuscript. PC analyzed the confocal images. JS edited the manuscript and was responsible for funding acquisition. JN conceived the study, designed the experiments, conducted the phosphoprotein array experiments, and was responsible for funding acquisition. All authors read and approved the final manuscript.

## Acknowledgements

The authors thank Dr. Tomáš Bárta (Department of Histology and Embryology, Faculty of Medicine, Masaryk University) for providing the NDF-2 cell line.

## Supplementary Information

**Fig. S1.**
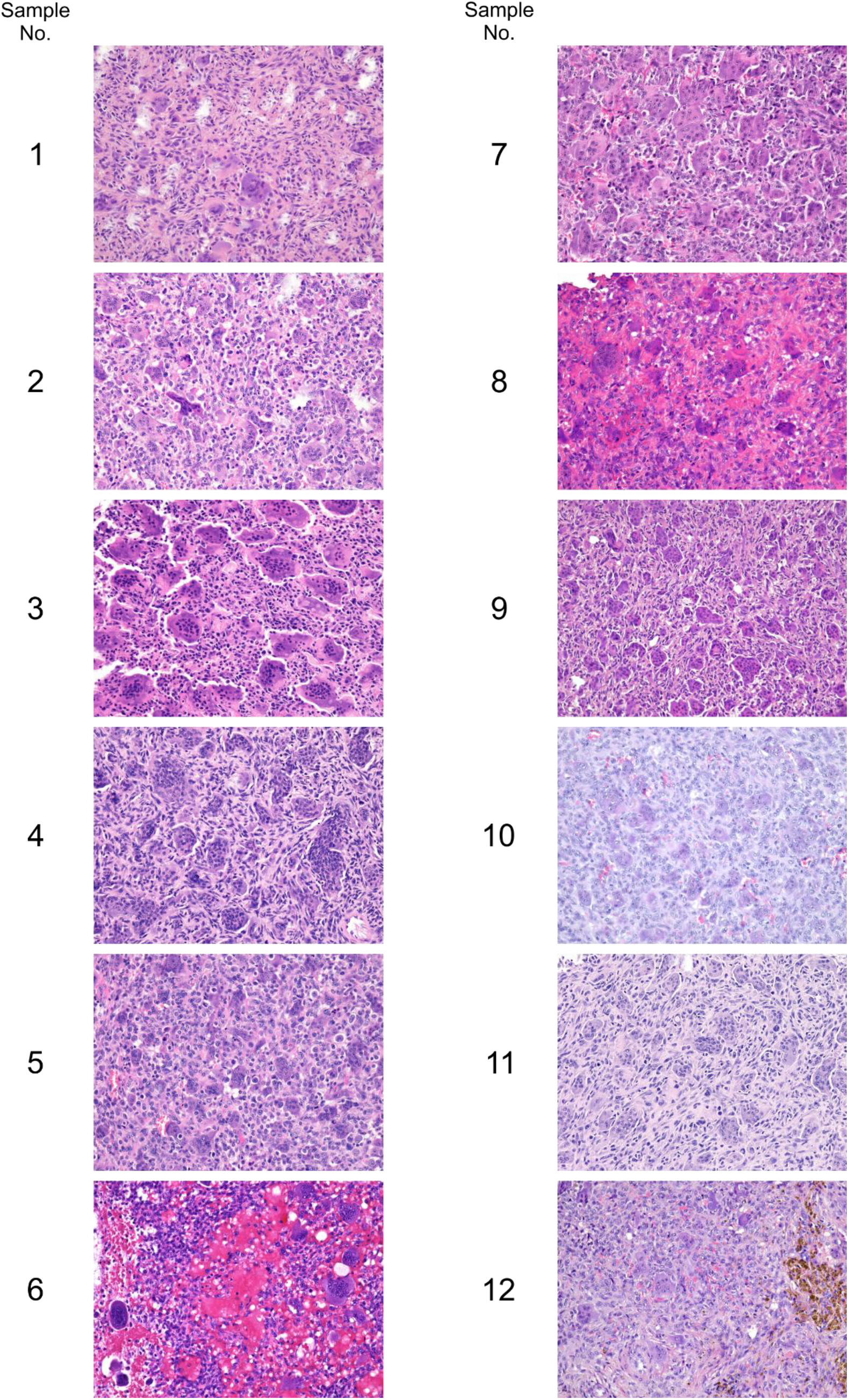
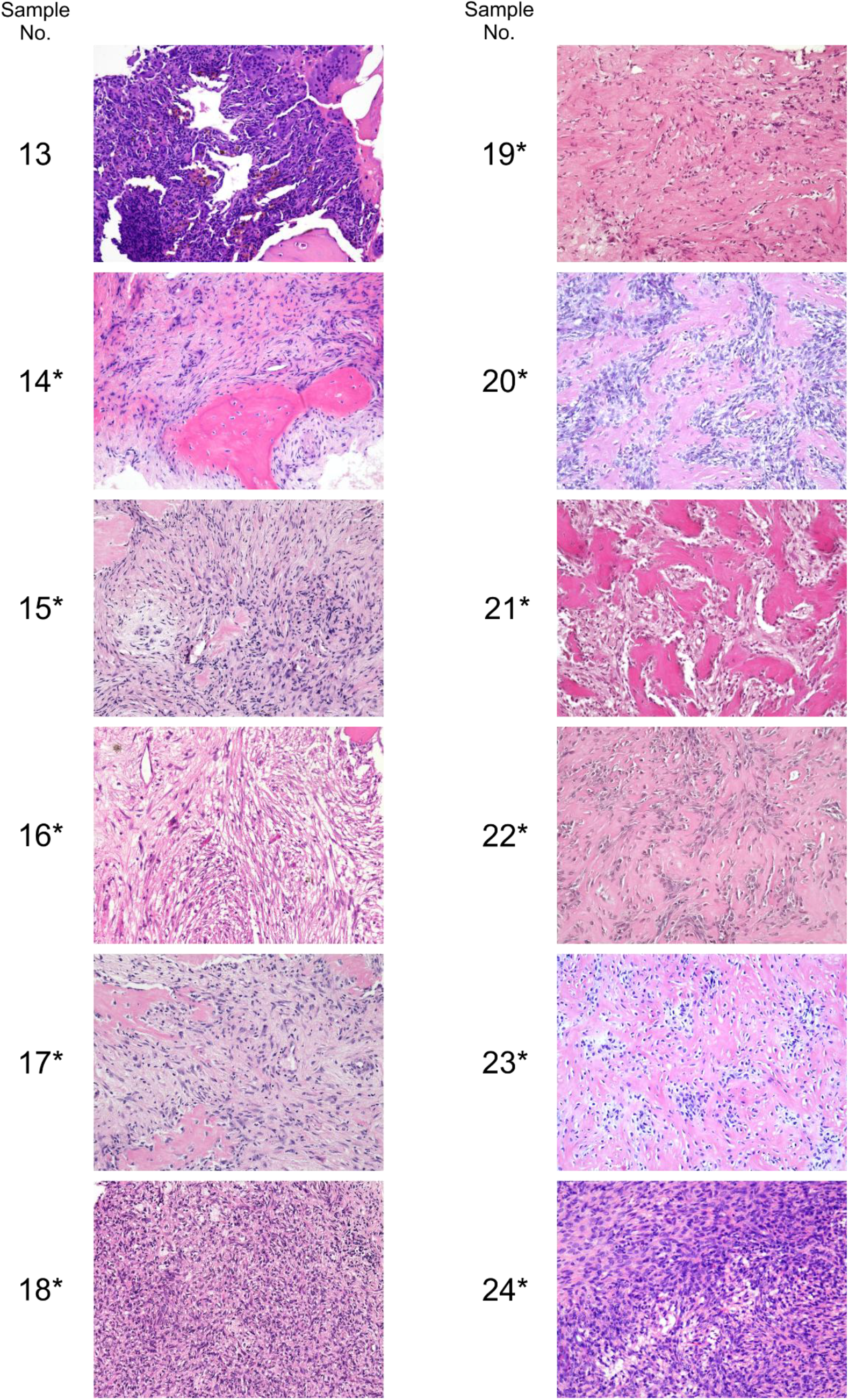
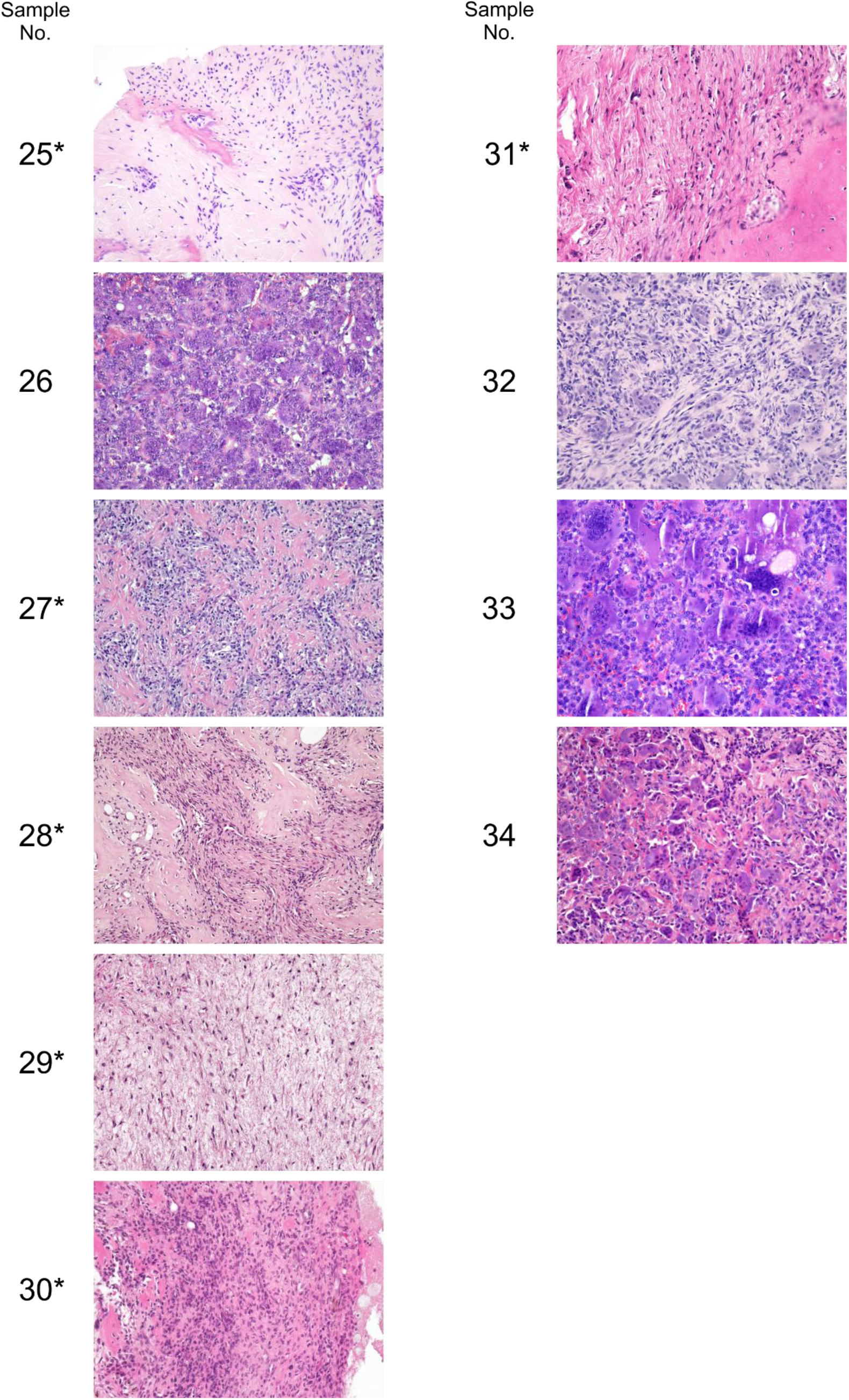
Histological analysis confirms a complete depletion of giant cells in denosumab-treated GCTB samples, whereas untreated samples contain both giant cells and neoplastic stromal cells. Denosumab-treated GCTB samples marked by asterisk (*).

**Fig. S2.**
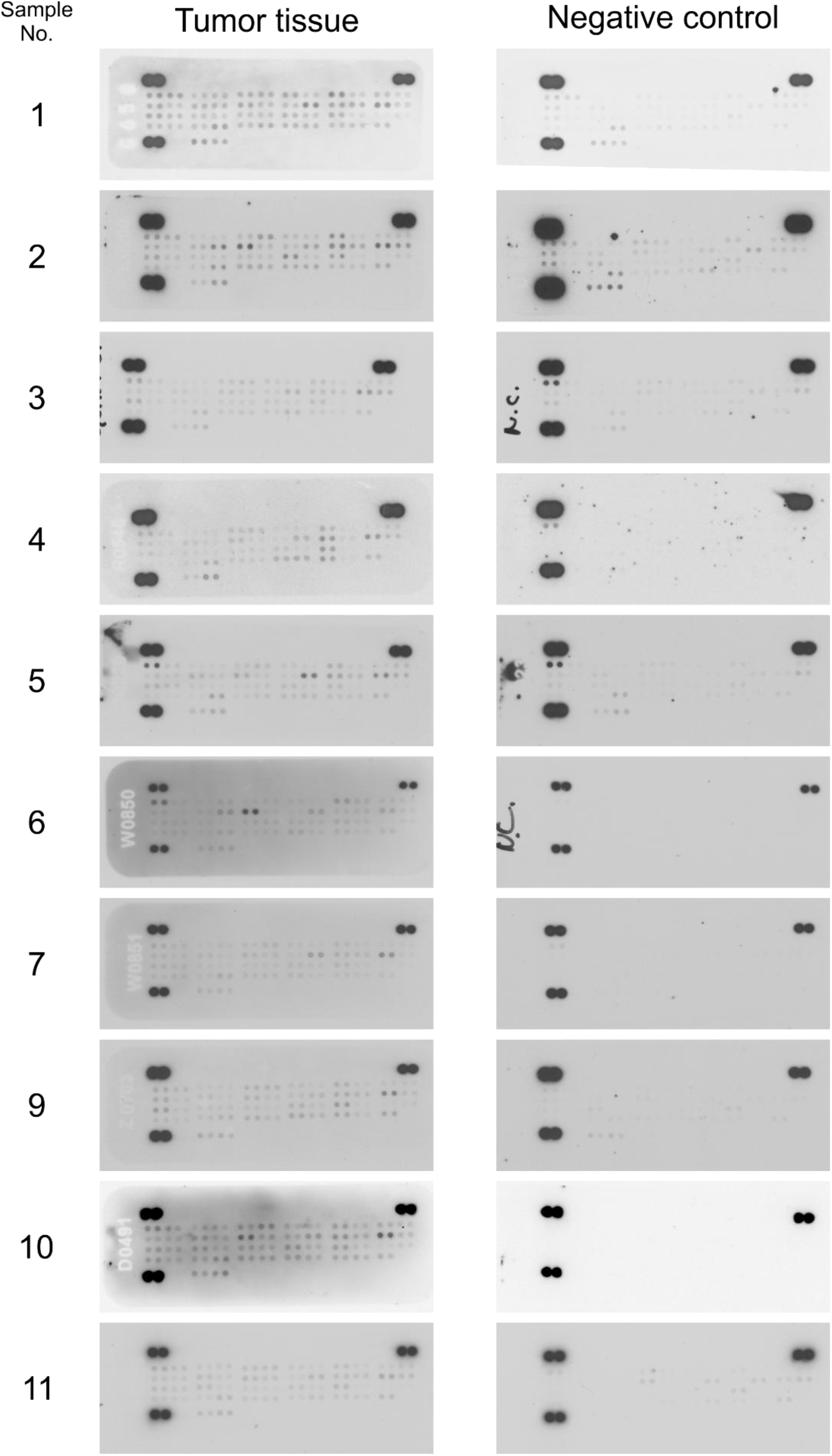
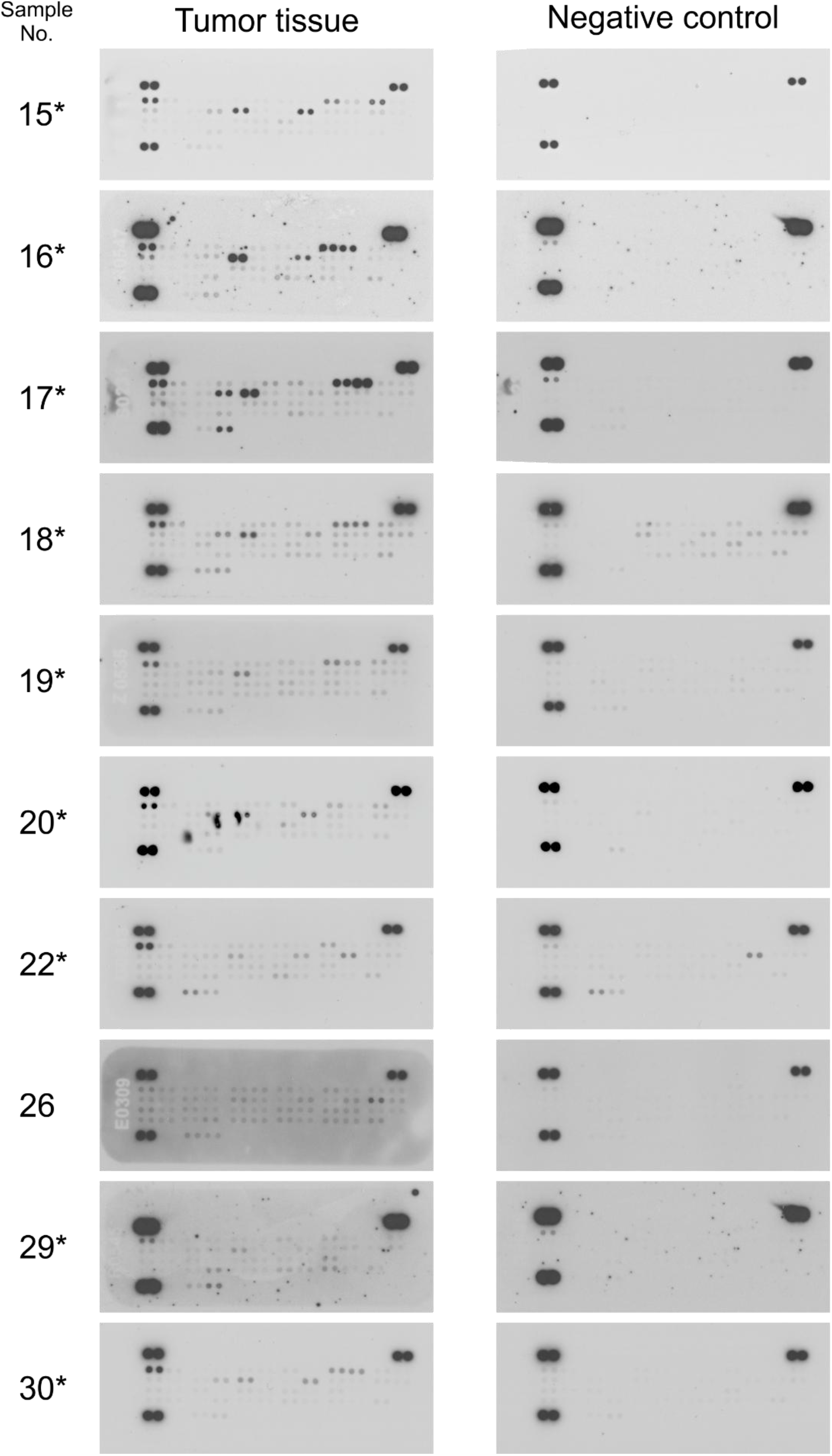
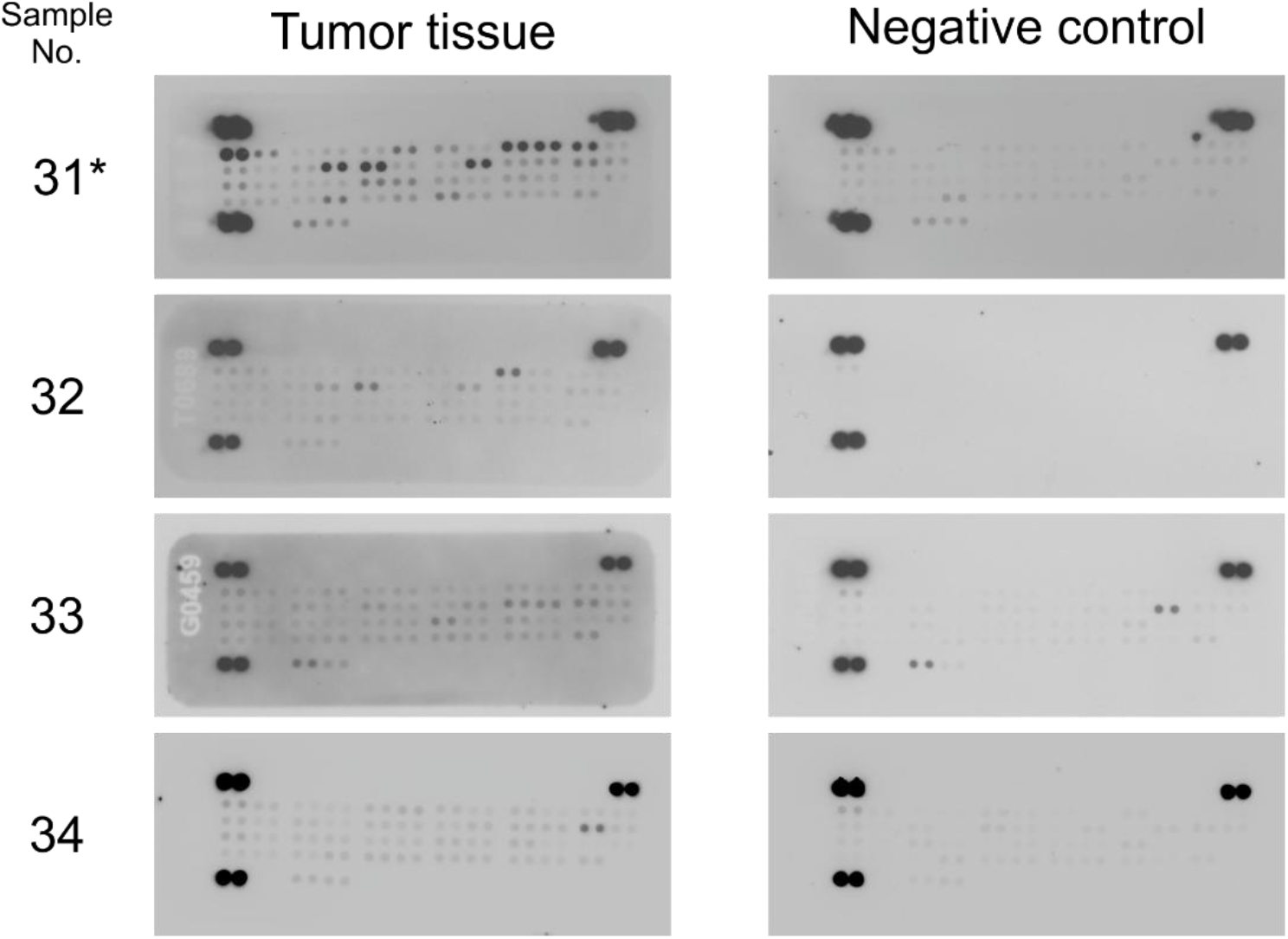
The images of the corresponding phospho-RTK arrays for all analyzed tissue samples, together with the respective negative controls, captured at the same exposure. Denosumab-treated GCTB samples marked by asterisk (*).

**Fig. S3.**
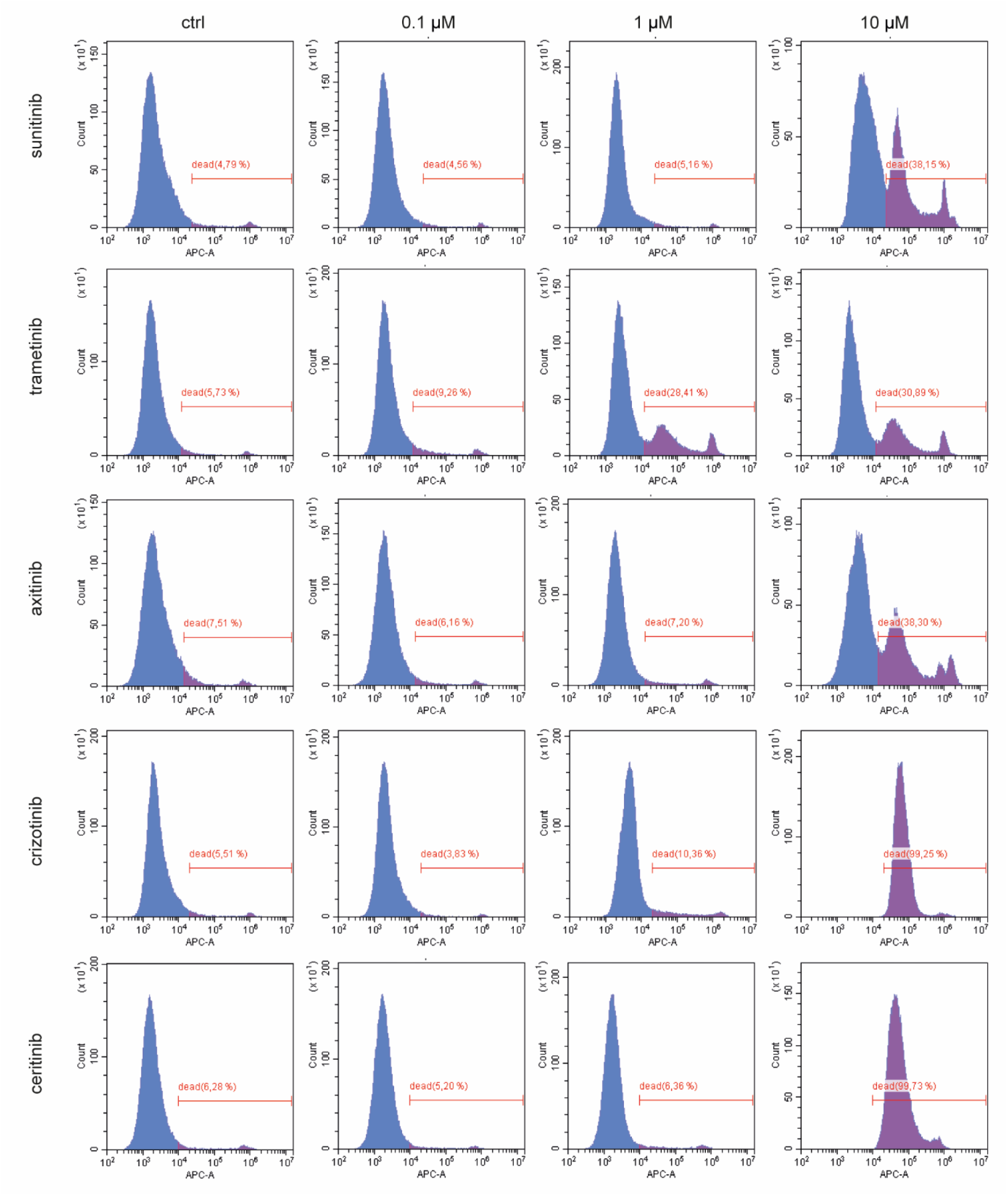
Flow cytometric analysis of dead cells after treatment with TKIs. The graphs show the gating strategy in one representative biological replicate of GCTB6 cell line treated with 0.1, 1 and 10 μM of sunitinib, trametinib, axitinib, crizotinib, and ceritinib.

**Fig. S4.**
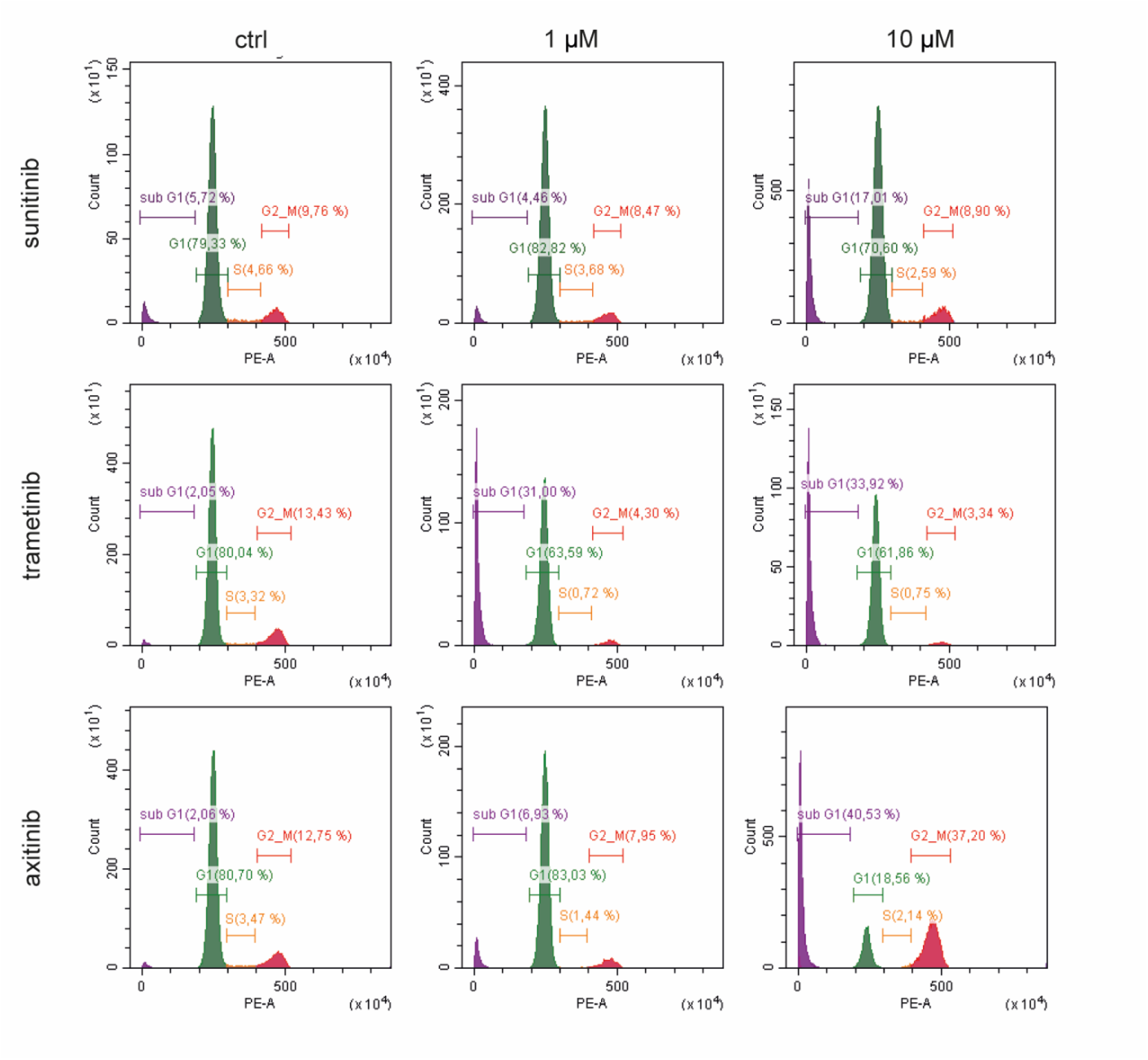
Flow cytometric analysis of changes in the cell cycle after treatment with TKIs. The graphs display the gating strategy in one representative biological replicate of GCTB6 cell line treated with 1 and 10 µM of sunitinib, trametinib, and axitinib.

